# Gene Regulatory Evolution During Speciation in a Songbird

**DOI:** 10.1101/029033

**Authors:** John H. Davidson, Christopher N. Balakrishnan

**Affiliations:** East Carolina University, Department of Biology, Greenville NC, USA

## Abstract

Over the last decade tremendous progress has been made towards a comparative understanding of gene regulatory evolution. However, we know little about how gene regulation evolves in birds, and how divergent genomes interact in their hybrids. Because of unique features of birds - female heterogamety, a highly conserved karyotype, and the slow evolution of reproductive incompatibilities - an understanding of regulatory evolution in birds is critical to a comprehensive understanding of regulatory evolution and its implications for speciation. Using a novel complement of analyses of replicated RNA-seq libraries, we demonstrate abundant divergence in gene expression between subspecies of zebra finches *Taeniopygia guttata.* By comparing parental populations and their F1 hybrids, we also show that gene misexpression is relatively rare, a pattern that may partially explain the slow buildup of postzygotic reproductive isolation observed in birds relative to other taxa. Although we expected that the action of genetic drift on the island-dwelling zebra finch subspecies would be manifested in a high rate of *trans* regulatory divergence, we found that most divergence was in *cis* regulation, following a pattern commonly observed in other taxa. Thus our study highlights both unique and shared features of avian regulatory evolution.

## INTRODUCTION

The study of gene expression in diverging species and their hybrids provides insights into the mechanisms of regulatory network evolution, adaptation and the origins of postzygotic reproductive isolation. Of particular interest to the process of speciation is gene misexpression, where expression in hybrids falls outside the range of variation observed in both parental populations (i.e., over- or under-dominance). Misexpression in hybrids may reflect Dobzhansky-Muller type incompatibilities and thus, can highlight the genetic changes underlying such incompatibilities (Michalak and Noor 2004). Over the last decade the comparative scope of gene regulatory evolution studies has expanded to include diverse study systems (e.g., Drosophila: Landry et al. 2005, Xenopus: Malone and Michalak 2008, whitefish: Renaut et al. 2009, yeast: Emerson et al. 2010, Busby et al. 2011, Schaefke et al. 2013). To date, however, no such study has been conducted in birds.

Birds display a number of traits that make them a particularly interesting target for studies of speciation genomics. First, they display female heterogamety where females are ZW and males are ZZ for their respective sex chromosomes. This feature allows for independent testing of sex chromosome-related features of speciation. For example, faster molecular evolution on the avian Z chromosome has been shown (Mank et al. 2010, Nam et al. 2010, Balakrishnan et al. 2013, Wright et al. 2015), following the pattern observed in many other taxa with heterogametic males (reviewed in Meisel and Connallon 2013). In terms of gene expression we may therefore expect to see faster expression evolution in Z-linked genes, and a tendency for Z-linked genes to be misexpressed in hybrids. Second, the evolution of reproductive isolation is protracted in birds relative to other taxa (Prager and Wilson 1975, Fitzpatrick 2004, Price 2008). Astoundingly, fully fertile hybrids have been documented from bird species that have diverged for up to 10 million years (Tubaro and Lijtmaer 2002, Lijtmaer et al. 2003, Price 2008, Arrieta et al. 2013). In many other taxa, studies of gene expression in have pointed to frequent misexpression in F1 hybrids (Landry et al. 2005, McManus et al. 2010, Malone and Michalak 2008, Renaut et al. 2009, Bell et al. 2013, Coolon et al. 2014). If gene misexpression in hybrids is reflective of the buildup of post-zygotic reproductive incompatibilties, then we may expect to see a reduced frequency of misexpression in bird species relative to other taxa of similar age.

The advent of RNA-seq technology has added a new dimension to the study of regulatory evolution as it is now possible to estimate the relative expression of alternative alleles across all expressed polymorphisms (McManus et al. 2010, Bell et al. 2013). Allelic imbalance, or allele specific expression, in F1 hybrids allows the further categorization of regulatory divergence into the contributions of *cis* and *trans* regulatory evolution and the interaction of the two. Most interspecific comparisons to date have found *cis* divergence to be more common (Landry et al. 2005, Tirosh et al. 2009, Goncalves et al. 2012, Graze et al. 2012). Others, however, have found a larger than expected contribution of *trans* divergence (McManus et al. 2010, Coolon et al. 2014). Based on comparisons between *Drosophila melanogaster* and *D. sechellia* McManus et al. (2010) hypothesized that demographic differences (increased drift) may drive this higher frequency of *trans* divergence. They posited that *trans* acting differences tend to generate intra-specific expression polymorphisms (Lemos et al. 2008, Wittkopp et al. 2008, Emerson et al. 2010) that might in turn be fixed by drift (Coolon et al. 2014).

Although there is extensive information about fertility and viability loss in hybrid birds (Tubaro and Lijtmaer 2002, Lijtmaer et al. 2003, Price 2008, Arrieta et al. 2013), to date there have been no studies of regulatory divergence in bird species and their hybrids. While zebra finches (*Taeniopygia guttata*) are an established model system for the neurobiology of song learning (Clayton et al. 2009), they also have great potential for mechanistic studies of speciation. In this study we examine regulatory divergence in two zebra finch subspecies, both of which are available in captivity and thus are readily amenable to experimental study. *Taeniopygia g. castanotis* and *T. g. guttata* inhabit mainland Australia and the Lesser Sunda islands of Southeast Asia, respectively. The Australian subspecies is broadly distributed across inland Australia whereas the Lesser Sundan subspecies (hereafter “Timor”) is found on the islands east of Wallace’s Line, a well-known biogeographic barrier (Wallace 1863, Huxley 1868). The subspecies appear to have diverged approximately one million years ago (Balakrishnan and Edwards 2009) when zebra finches colonized the Lesser Sunda islands from Australia (Mayr 1944). The two subspecies are reciprocally monophyletic for mtDNA alleles (Newhouse and Balakrishnan in review) but not for any nuclear markers surveyed to date (Balakrishnan and Edwards 2009). Patterns of genetic variability suggest the colonization of the islands involved a substantial population bottleneck that is reflected in much reduced genetic diversity among island birds. Here we broadly describe patterns of expression divergence between zebra finch subspecies and in doing so, test whether genetic drift resulting from a historical bottleneck has impacted patterns of regulatory evolution in zebra finches.

## RESULTS

### Differential Expression Between Subspecies

Nine RNA-seq libraries, derived from RNA extracted from the brains of three Australian, three Timor and three F1 hybrid zebra finches, yielded over 30 million reads per sample (Table 1). Using *bwa mem* (Li et al. 2009) we were able to map over 85% of our reads to a version of the zebra finch genome that had been masked of SNPs fixed for alternative alleles in Australian and Timor finches. Across all nine libraries, we detected 16,689 out of 18,618 (89.6%) Ensembl-annotated genes with at least one read in one library. The brain-expressed transcripts we detected were a non-random representation of the genome as a whole. For example, GO categories “structural constituent of cytoskeleton” (000520), “G-protein coupled receptor signaling pathway” (0007186), and “intermediate filament” (0005882) were all strongly underrepresented among our detected transcripts (Fisher’s Exact Test *p* < 0.05). Although olfactory receptors are expressed in the brain, expression is highly localized, thus genes annotated with related GO annotations (0004984, 0050911) were also significantly under-represented in our whole brain transcriptome. The dataset was also enriched for a number of GO categories, mostly having to do with cellular components (e.g., cytoplasm (0005737), nucleus (0005634), mitochondrion (0005739); see also Supplementary Material). Functional components that were enriched included protein binding (000551), hydrolase activity (0016787) and transferase activity (0016740).

**Table 1.** Total number of reads after quality trimming and proportion mapped to the reference genome before and after masking polymorphic sites.

|  | Trimmed Reads | tophat2 initial | tophat2 masked | bwa mem Initial | bwa mem Masked |
| --- | --- | --- | --- | --- | --- |
| **Australian** |  |  |  |  |  |
| **Library 1** | 31,726,619 | 0.684 | 0.472 | 0.9 | 0.885 |
| **Library 2** | 33,049,620 | 0.669 | 0.455 | 0.876 | 0.871 |
| **Library 3** | 32,844,330 | 0.669 | 0.455 | 0.874 | 0.869 |
| **average** |  | 0.674 | 0.461 | 0.883 | 0.875 |
| **Timor** |  |  |  |  |  |
| **Library 1** | 33,115,938 | 0.643 | 0.464 | 0.883 | 0.879 |
| **Library 2** | 34,328,721 | 0.645 | 0.466 | 0.883 | 0.879 |
| **Library 3** | 33,467,969 | 0.621 | 0.443 | 0.864 | 0.86 |
| **average** |  | 0.636 | 0.458 | 0.877 | 0.873 |

After filtering for variance outliers under default settings in DE-Seq2 (Anders and Huber 2010), 13,904 genes were tested for differential expression. Of these, 913 genes (6.6%) were differentially expressed between Australian and Timor zebra finches (*p* < 0.05). All *p*-values from DE-Seq2 analyses were adjusted for multiple testing (Benjamini and Hochberg 1995). Of the differentially expressed genes, 51.5% were up-regulated in Australian finches and the remaining 48.5% were up-regulated in Timor zebra finches. Thus the distribution of fold changes across all genes was centered around zero with no tendency of genes to be up-regulated in one population versus the other. Among the differentially expressed genes, those with roles in oxidation-reduction process (0055114, 44 genes) and oxidoreductase activity (0016491, 36 genes) were significantly enriched. In both of these GO categories just over 50% (52.3, 52.7%) of the transcripts were more highly expressed in Australian Zebra Finches. Genes with annotated roles in protein binding, sequence-specific DNA binding, and transcription factor activity were under-represented, suggesting relatively conserved expression of genes in these categories. No KEGG pathways were significantly enriched or underrepresented after correcting for multiple testing. However MAPK-related genes (gga04010) were relatively conserved in expression with only one differentially expressed gene in this pathway (expected = 8, Fisher’s Exact Test, *p* = 0.17). Oxidative Phosphorylation-related genes (gga00190) were slightly enriched with 10 such genes differentially expressed (expected = 4, Fisher’s Exact Test *p* = 0.17). Eight out of the ten oxidative-phosphorylation-related genes were expressed more highly in Timor finches relative to Australian zebra finches (binomial test, *p* = 0.04).

### Expression Divergence on the Sex Chromosome

We tested for elevated regulatory divergence on genes of the Z chromosome by comparing the variance in fold-changes across the Z relative to chromosome 4, the chromosome most similar in gene content (number of genes). If Z chromosome genes were diverging more rapidly in terms of expression, we would expect a larger variance in fold change. However, we found no significant difference in variance among chromosomes (F = 1.03, *p* = 0.72). We also found no enrichment of genes on the Z among those that were differentially expressed between subspecies. Whereas Z linked genes comprise 4.6% of the detected genes in our dataset (793/13,904), 3.6% (33) Z-linked genes were differentially expressed. This difference was not statistically significant (χ^2^ = 2.02, *p* = 0.15). Thus, Z-linked genes are not significantly over- or under-represented among the differentially expressed genes.

### Inheritance of Gene Expression

We also classified the mode of inheritance of expression profiles in hybrid birds relative to the parental subspecies. We successfully classified inheritance for 847 differentially expressed genes. In contrast to many previous studies in non-avian taxa, we found only five genes (0.5%) with significant evidence of misexpression in hybrids (p < 0.05, Figure 1B, Figure 2): AP3B2, POMC, WNT7A, EFCAB2, AKR1b (gene family member). At a less stringent significance threshold of *p <* 0.10, only one additional gene, TFIP11, can be classified as misexpressed (Figure 2). Instead, the vast majority of genes, 631 in total (74.5%), showed an additive inheritance pattern. Another 211 genes showed dominance with 169 dominant in the Timor zebra finches over their Australian counterparts, and only 42 showed the reverse (χ^2^ = 70.6, *p* < 0.0001).

**Figure 1.**
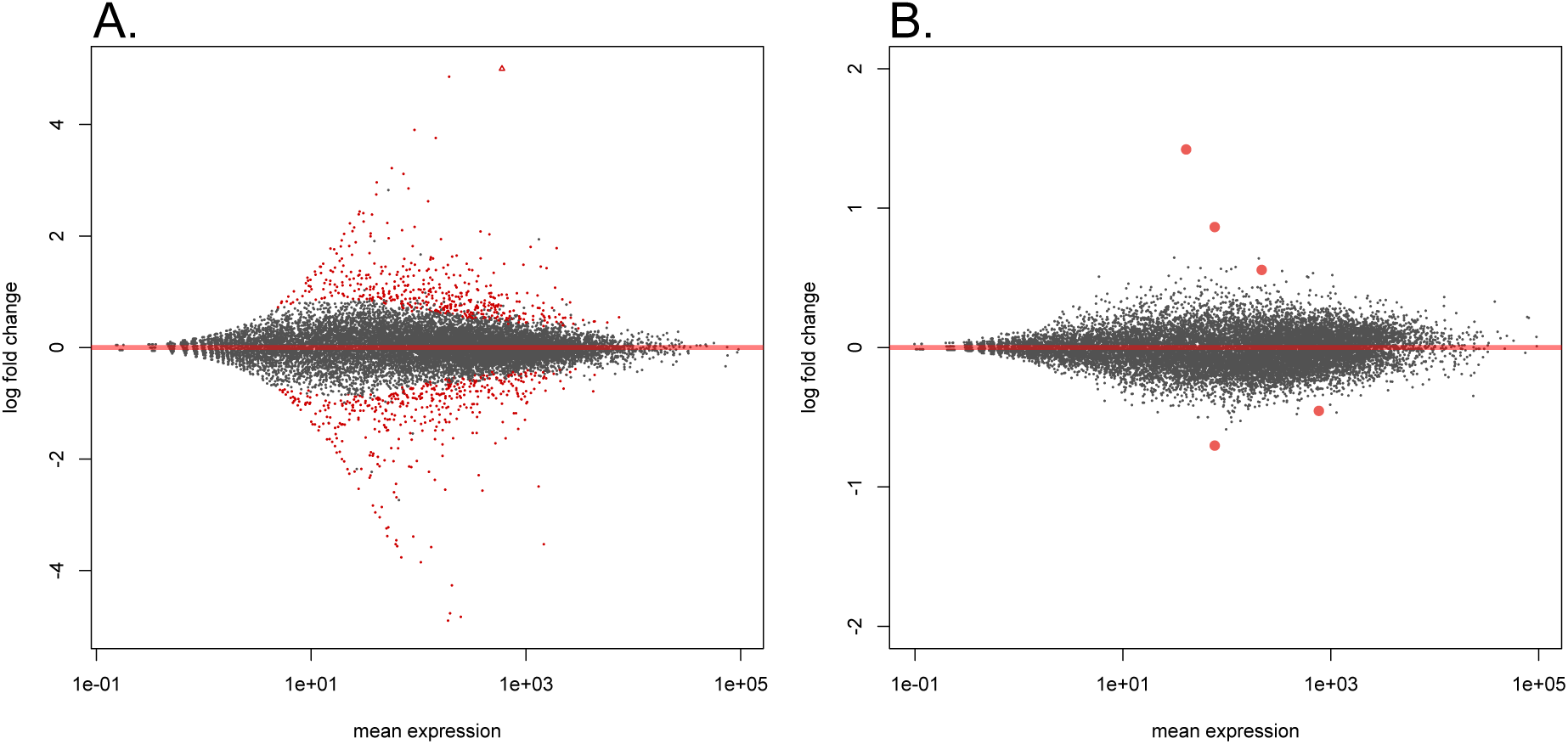
MA Plot (expression level versus log fold change) of differential expression for two contrasts: A) Australian versus Timor zebra finches and B) Parental subspecies versus their hybrids. Points in red are significant at *p* < 0.05 (adjusted for multiple testing).

**Figure 2.**
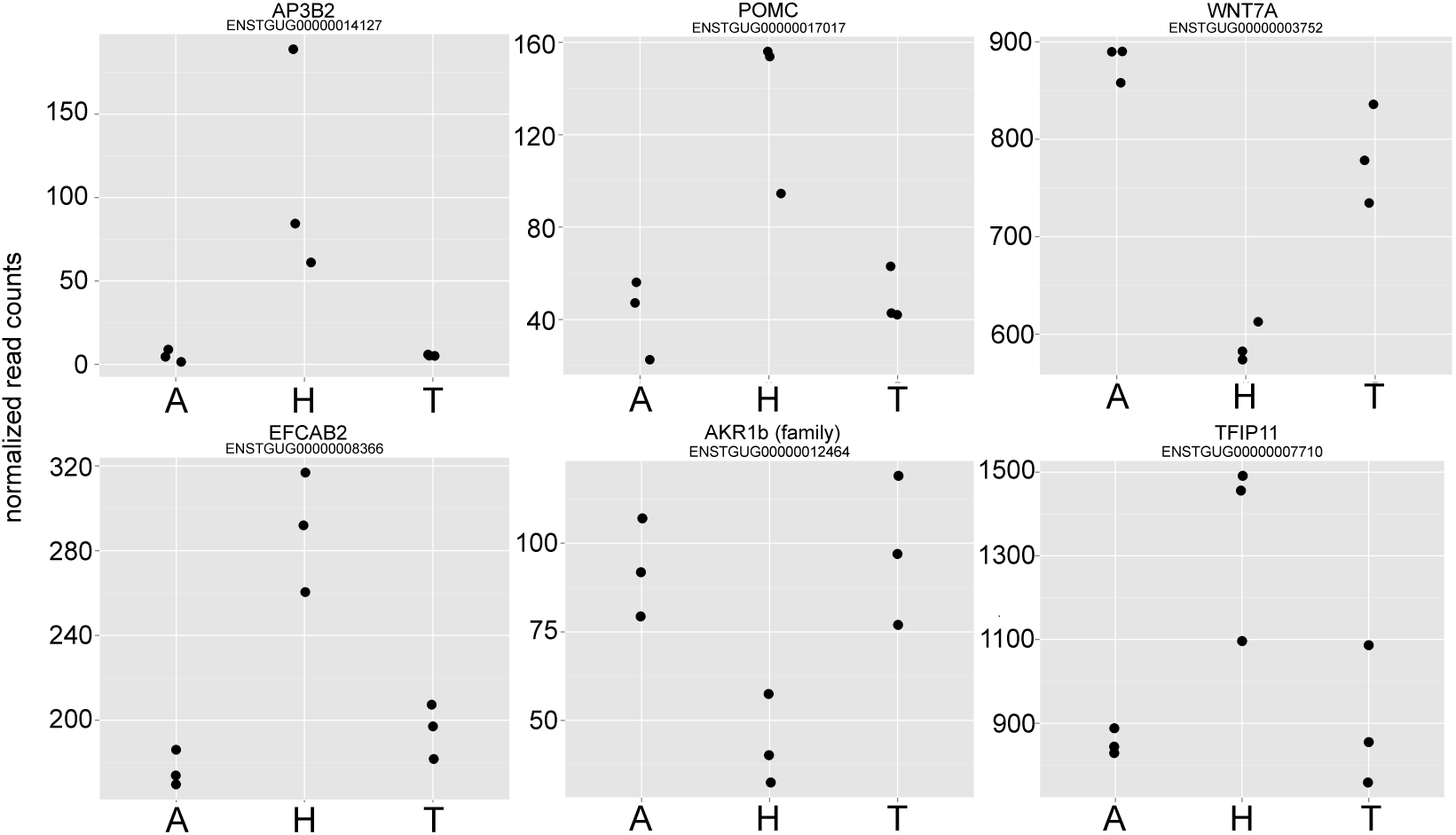
Six misexpressed genes in hybrid zebra finches. Statistics are based on differential expression comparison of the two zebra finch subspecies (n=6) versus their hybrids (n=3) (Figure 1B). Five of these genes are significant at adjusted *p* < 0.05 and the sixth (TFIP11) is significant at adjusted *p* < 0.1.

### Mode of Regulatory Divergence

Biased allelic expression in F1 hybrids reflects *cis* regulatory divergence between parents, since alleles inherited from each parent are exposed to the same *trans* regulatory environment (Wittkopp et al. 2004). Because we had replicated RNA-seq libraries we were able to use statistical software tailored for RNA-seq, thus incorporating observed variance profiles within and among genes to test for both allelic bias in hybrids and *trans* divergence. *Trans* divergence is identified as differences in allelic ratio in hybrids versus the ratio of genic expression among parents (Wittkopp et al. 2004, see Methods). We assessed patterns of allelic expression in 23,838 SNPs whose genotype was ascertained for all nine samples. This set of SNPs included only sites for which the two subspecies were fixed for alternative alleles, allowing unambiguous determination of ancestry in the F1 hybrids.

Of the SNPs we tested for allele-specific expression, only 6,634 (28%) mapped within annotated genes (including exons, UTR, introns). The majority of SNPs, 12,129 (50.1%) in total, mapped outside of gene models but within 5kb downstream of known genes, possibly representing unannotated UTR regions. Because the annotation of the zebra finch genome is incomplete, gene associations of these and other noncoding SNPs are uncertain. Thus, we conducted our analysis at the level of individual sites rather than genes (Bell et al 2013), recognizing that some SNPs will be non-independent because they are associated with the same gene.

If genetic drift has led to an accumulation of deleterious alleles in Timor zebra finches, one pattern we might expect to see is a tendency towards higher expression of Australian alleles. In general, however, the two alternative alleles were expressed equally in hybrid birds (22,658/23,838 SNPs, 95%). Of the remaining sites, we found significant evidence of biased allelic expression, and thus *cis* divergence between parents, in 253 SNPs (1% of sites, Figure 3). Two hundred and twenty-five of the 253 SNPs were putatively associated with 155 annotated genes (in the UTR, intron, exon or within 5kb up or downstream) and the remaining 28 SNPs were intergenic. Even among the sites where we observed biased expression in hybrids, the average log2 fold change was zero. Thus there was no bias in terms of which allele (Timor or Australian-derived) was more highly expressed.

**Figure 3.**
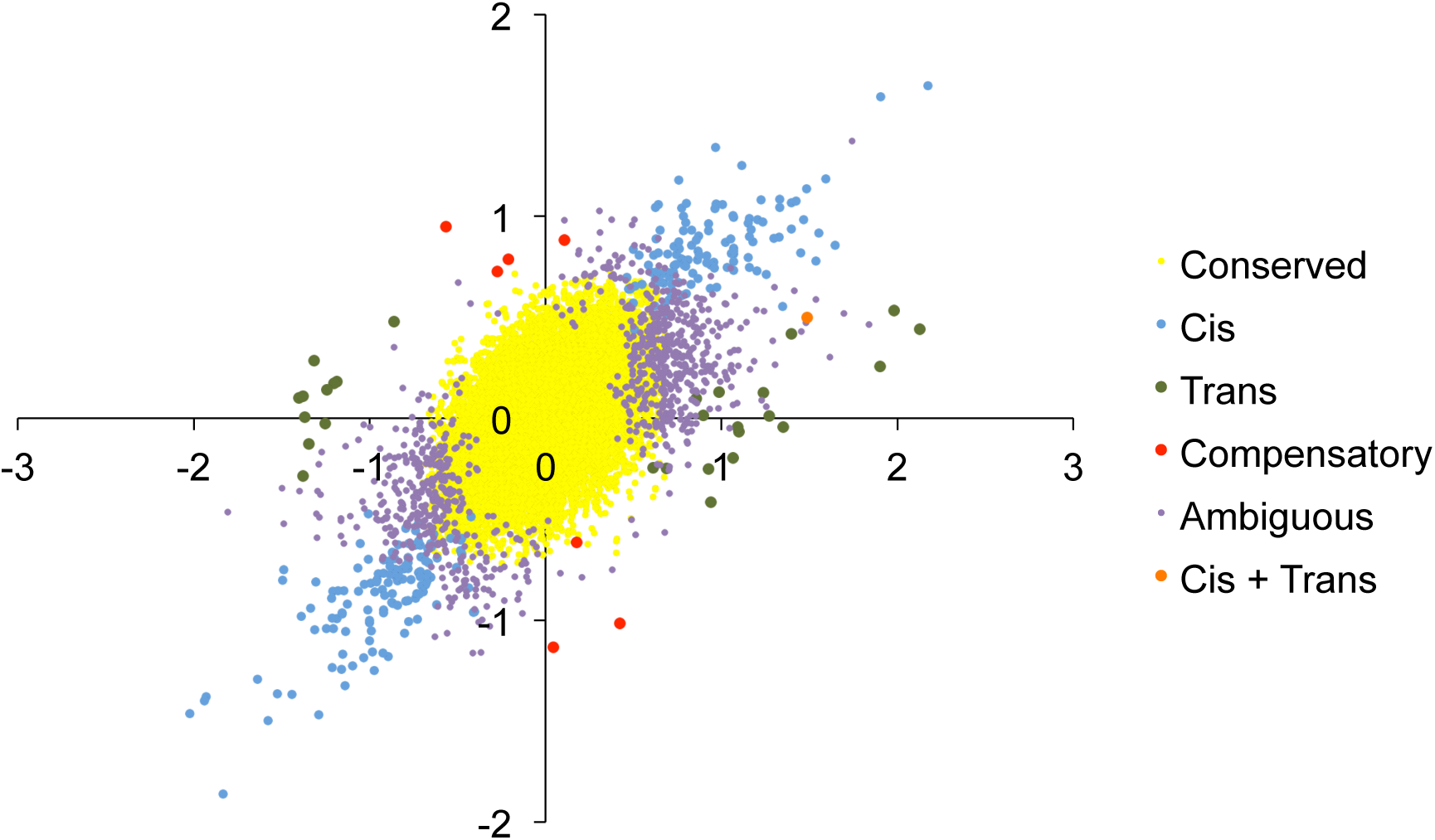
Categorization of regulatory divergence modes based on patterns of allele-specific expression in hybrids and subspecific divergence. Most loci showed conserved expression (yellow), however, among those that show significant evidence of evolution changes in *cis* regulation were most common (light blue).

We combined information from allelic bias in F1 hybrids with information on expression differences between parents and a test for *trans* divergence to further categorize regulatory divergence into subcategories (Table 2, Figure 3). Another 53 sites showed significant *trans* divergence. Of these, 28 sites putatively representing 17 genes showed *trans* only divergence. Seven sites showed evidence of compensatory evolution, where there was *cis* and *trans* regulatory divergence, but no net divergence in overall expression between subspecies. Only one site showed significant divergence in expression between subspecies, significant *cis* divergence, and significant *trans* divergence. This represents a lone case of *cis* and *trans* regulatory divergence acting together to cause expression divergence between subspecies. Eight hundred and ninety-one sites revealed an ambiguous pattern and could not clearly be categorized in the their mode of divergence.

**Table 2.** Overview of classification scheme for categorizing patterns of regulatory divergence. “Yes” or “no” refers to a significant statistical test as defined in the methods.

| Mode | Parental Divergence | ASE in hybrids | TransTest |
| --- | --- | --- | --- |
| ***cis* only** | yes | yes | no |
| ***trans* only** | yes | no | yes |
| ***cis* x *trans*, *cis* + *trans*** | yes | yes | yes |
| **compensatory** | no | yes | no |
| **conserved** | no | no | no |

## DISCUSSION

In this study we have broadly characterized the regulatory divergence of brain-expressed transcripts in two zebra finch subspecies that have been geographically isolated for around one million years (Balakrishnan and Edwards 2009). We find evidence of abundant expression divergence between the two populations, with over 900 genes showing differential expression. Among these genes, those involved in oxidation-reduction and metabolic processes were significantly enriched. The divergence in genes associated with metabolism, including a mild enrichment of oxidative phosphorylation-related genes, could be related to ecological adaptation to different habitats in inland Australia versus the Lesser Sunda islands. Alternatively, it is possible that differences in expression are the result of short-term adaptation to captivity in the Australian subspecies (< 200 years, Zann 1996). The tendency for oxidative-phosphorylation genes to be more highly expressed in Timor zebra finches (8 out of 10 differentially expressed genes) supports these adaptive scenarios.

Unlike many previous studies (Landry et al. 2005, McManus et al. 2010, Malone and Michalak 2008, Renaut et al. 2009, Bell et al. 2013, Coolon et al. 2014), we found that misexpression was rare among the genes we measured in hybrid zebra finches. Even though we find subspecies differences in expression of genes related to cellular energetics, we don’t find any evidence of misexpression of mitochondrial or other oxidative phosphorylation-related genes in hybrids. Mito-nuclear interactions are known to contribute to genomic incompatibilities in certain taxa (e.g., Burton et al. 2006, Ellison and Burton 2006, Ellison et al. 2008) and have recently been suggested to be particularly likely candidates as “speciation genes” (Burton and Barreto 2012, Hill 2015). In zebra finches, we know that mtDNA alleles are differentiated between the two species (Newhouse and Balakrishnan in review), yet we don’t find evidence of resultant misregulation of mitochondrial genes.

A number of factors may contribute to the low levels of misexpression in zebra finch F1 hybrids. Although the two zebra finch subspecies are relatively divergent, they are young relative to some species pairs previously tested for misexpression (e.g. *simulans x D. melanogaster,* 2.5 million years, Landry et al. 2005, *Sacchromyces cerevisiae* x *S. paradoxus*, 5 million years, Tirosh et al. 2009). On the other hand, even crosses between relatively young mouse subspecies (< 0.5 MY) exhibit hybrid sterility and misexpression (Mack et al. in review). Given that there is no evidence of reproductive incompatibility between the zebra finch subspecies and that post-zygotic isolation takes a relatively long time to evolve in birds (Prager and Wilson 1975), our results suggest that misexpression may accumulate after the origin of reproductive incompatibilities or may directly contribute to the origin of incompatibilities.

Its also important to note that in this study we examined only brain tissue. If patterns of expression in the brain are relatively conserved, that could contribute to reduced levels of misexpression observed here. Unlike many studies of *Drosopila,* Graze et al. (2012) examined gene expression in dissected heads (rather than whole bodies) of *D. melanogaster* x *D. simulans* hybrids, and they too found limited evidence of misregulation (~30 genes). Our brain transcriptome, however, included the majority of annotated genes, suggesting that if misexpression were pervasive, we would detect it even in brain tissue. We have also examined only F1 hybrids. Examination of F2 or backcrossed individuals remains an important goal for the future as second generation crosses allow recombination among parental genomes, potentially exposing deleterious allelic combinations. For example, studies of wild whitefish populations showed relatively low frequency of misexpression in F1 hybrids (9% of genes) but abundant misexpression in backcrosses (54% of genes) (Renaut et al 2009). The importance of second generation hybrids is particularly true for incompatibilities derived from mito-nuclear interactions where such crosses can pair a mitochondrial genome from one genetic background with the nuclear genetic background of the other.

Possibly the most interesting implication of our findings is if the lack of misexpression in zebra finch hybrids is related to the long-known pattern of slow post-zygotic reproductive isolation evolution in birds (Prager and Wilson 1975). Thorough testing of this hypothesis, however, will require examination of additional tissues and species pairs. A number of hypotheses for the slow buildup of incompatibilities in birds have been proposed. Fitzpatrick (2004) favored a hypothesis of slower regulatory evolution in birds than in mammals. Our results don’t support this hypothesis as we observed substantial amounts regulatory divergence between subspecies, yet little evidence of misregulation in hybrids. Another hypothesis is that differences in dosage compensation and sex chromosome systems are responsible (Fitzpatrick 2004). The idea is that in mammals, X inactivation in females causes deleterious recessive mutations on the X to be exposed in both sexes, whereas in birds, males express their diploid Z chromosome genotype. In this study we tested only males, thus it possible that an examination of females would expose more widespread misregulation of sex-linked genes and their interaction partners. A novel hypothesis is that the slow rate of evolution of post-zygotic incompatibility is due to the stability of the avian karyotype (e.g., Griffin et al. 2007). If changes in genomic architecture (e.g., inter chromosomal rearrangements) contribute to regulatory divergence and hence, to genetic incompatibilities, perhaps in birds it simply takes longer for such changes to accrue.

The small number of a misregulated genes does not necessarily imply that misregulation is unimportant in this system. One gene that was clearly misexpressed is proopiomelanocortin (POMC, Figure 2). POMC is notable as a gene with multifaceted roles in pigmentation and social behavior. Pigmentation patterns clearly differ among subspecies and aspects of social behavior may vary as well. It will be of interest to determine whether regulatory changes in POMC or any other misexpressed gene contributes to any phenotypic differences in the zebra finch subspecies, and whether the misexpression that does occur causes aberrant phenotypes in hybrid offspring.

Studies of sex chromosome evolution have revealed that like the mammalian X, the avian Z is evolving rapidly relative to autosomes. This pattern has been attributed primarily to genetic drift (Mank et al. 2010), and is further modulated by variation in the strength of sexual selection (Wright et al. 2015). Surprisingly, we found no unusual patterns of expression divergence for Z-linked genes. At the nucleotide level, the Z chromosome has been demonstrated to be evolving relatively quickly in estrildid finches (Balakrishnan et al. 2013), the group to which zebra finches belong. These changes must not be influencing gene expression, or compensatory substitutions may be mitigating the consequences of deleterious changes. However, we were able to detect only a handful of compensatory changes and none of these were on the Z chromosome.

The Timor zebra finch subspecies has undergone a severe bottleneck in colonizing the Lesser Sunda islands as evidenced by dramatically reduced neutral genetic variation (Balakrishnan & Edwards 2009). That pattern as well as patterns of trait evolution led Balakrishnan and Edwards (2009) to conclude that a founder effect likely played a role in zebra finch speciation. Under strong genetic drift, we expected to see relatively abundant *trans* regulatory divergence if many deleterious alleles were fixed in the island populations (McManus et al 2010). Such a pattern was observed in comparisons of *D. melanogaster* and *D. sechellia,* the latter of which also is an island form (McManus et al 2010). However in zebra finches, as in a number of recent studies (Landry et al. 2005, Tirosh et al. 2009, Goncalves et al. 2012, Graze et al. 2012), we find that most of the regulatory divergence was *cis* acting. We also predicted that under drift, deleterious mutations would accumulate and impair normal levels of gene expression. In this case, Timor zebra finch alleles would show a tendency to be underexpressed relative the Australian allele in heterozygous hybrids. Again though, we do not see this pattern. Taken together, we don’t find compelling evidence of bottleneck-induced drift influencing patterns of gene expression.

Two components of our analysis showed asymmetry with respect to patterns of expression in the two subspecies. First, the Timor expression pattern tended to be dominant over that of the Australian subspecies (169/211 genes) and second, oxidative-phosporylation genes tended to be more highly expressed in Timor zebra finches (8/10 differentially expressed genes). Genetic drift due to bottlenecks during domestication of Australian zebra finches may provide an explanation for the former pattern, whereas adaptation to captivity may explain the latter pattern. Subtle genetic divergence between wild and domesticated Australian zebra finches has been described, but these populations remain relatively outbred and polymorphic (Forstmeier et al 2010, Newhouse and Balakrishnan in review).

Finally, on a technical note, we presented a simple framework by which both differential expression and allelic specific expression (and thus, the contributions of *cis* and *trans* divergence) can be assessed in a consistent statistical framework using replicated experiments and statistics tailored for RNA-seq data. Previous RNA-seq based studies of regulatory divergence have estimated allelic bias using binomial tests, which do not take into account sample variance and the expected and observed pattern of variance in RNA-seq data. Here we have used a single software package, DE-Seq2 to test for divergence in gene expression, allelic expression and the interaction of the two, or *trans* divergence. Specifically, we used an interaction term in a general linear model to test for *trans* divergence. We suggest that this approach using DE-Seq2 or similar software, paired with phylogenetic studies of regulatory divergence (e.g., Coolon et al 2014) will continue the progress toward a broad, comparative understanding of regulatory evolution.

## MATERIALS AND METHODS

### RNA Extraction, Library Preparation and Sequencing

Birds were housed in captivity at the Institute for Genomic Biology at the University of Illinois at Urbana-Champaign. Three male birds were sampled from each of three populations, Australian (*T guttata castanotis),* Timor (*T guttata* guttata) and hybrid finches. All of the hybrid birds studied were the result of crosses between female Australian zebra finches and Timor males. This crossing directionality was chosen because female Australian zebra finches breed more readily in captivity than do female Timor finches. In order to control for environmental influences on gene expression, each individual bird was placed in an acoustic isolation chamber the night before they were to be sacrificed. To avoid pharmacological influences on gene expression, birds were then euthanized by decapitation. Tissues were dissected and then snap-frozen on dry ice. All animal protocols were approved by the University of Illinois IACUC. All procedures subsequent to dissections were carried out at East Carolina University.

Whole brain tissue was homogenized in Tri-Reagent (Molecular Research Company) for RNA purification and total RNA was extracted following manufacturer’s instructions. Total RNA was then DNase treated (Qiagen) to remove any genomic DNA contamination and the resulting RNA was further purified using RNeasy columns (Qiagen). Purified total RNA was assessed for quality using an Agilent Bioanalyzer. Library preparation and sequencing were done at the University of Illinois Roy J. Carver Biotechnology Center. Library preparation used Illumina TruSeq RNA Sample Prep Kit and manufacturer’s protocols. RNA Sequencing was performed in a single lane of an Illumina HiSeq 2000 using a TruSeq SBS sequencing kit version 3 producing single end 100 base pair reads which were analyzed with Casava 1.8.2. Reads were adapter and quality-trimmed using *Trim Galore!* (Kreuger 2015), a wrapper script that uses *cutadapt* (Marcel 2011) to trim reads.

### Read Mapping, Expression Measurement, and Differential Expression Testing

We expected that reads from Australian zebra finches would map to the reference genome (v3.2.74) at a higher rate than would Timor zebra finches because the genome was derived from the Australian subspecies. We observed such biases in preliminary analyses using *bwa aln* (Li et al. 2009) and *tophat2* (Kim et al. 2013; Table 1). We observed little bias, however, when we mapped we mapped reads to the genome using *bwa mem* (Li et al. 2009) under default settings (Table 1). Therefore, we used this read aligner for subsequent analyses. Despite the apparently consistent mapping of reads at the whole genome scale, mapping bias at specific loci that are divergent in sequence could still preclude accurate expression measurements. To avoid this we masked sites in the genome with fixed differences between subspecies. To accomplish this, we identified polymorphisms in the dataset using *samtools mpileup* (Li et al. 2009) and called SNPs using *bcftools* (Li et al. 2009). Fixed differences were then identified using *SNPSift* (Cingolani et al. 2012), filtering the VCF (variant call format) file generated by *bcftools* for sites that were homozygous for the reference allele in the three Australian birds and homozygous for the alternative allele in the Timor birds. These sites were then masked in the reference genome using *bedtools* (Quinlan and Hall 2010). Following masking we re-mapped reads to the masked genome again using *bwa mem.* The proportion of mapped reads dropped by about 1 percent after masking fixed differences (Table 1) but we used the masked mapping to avoid any potential bias in downstream analyses.

We quantified gene expression relative to Ensembl-annotated gene models (Ensembl v73). For each gene, we counted the number of overlapping reads using ht-seq (Anders et al. 2014). We then used DE-Seq2 (Anders and Huber 2010, Love et al. 2014) to normalize read counts per library and to test for differential expression. We conducted four pairwise tests to categorize genes as differentially expressed among species, but also to categorize inheritance as dominant/recessive, additive, or over/under-dominant. Together, we consider the latter two categories as being “misexpressed”. The pairwise tests were: Australian (n=3) vs. Timor (n=3), hybrids (n=3) versus parentals (n=6), hybrids (n=3) versus Australian (n=3) and hybrids (n=3) versus Timor (n=3). Inheritance was considered additive if expression in hybrids was intermediate to the two parentals but not significantly different from either parental subspecies. If hybrid expression was intermediate, but expression in hybrids was significantly different from one parent but not the other, inheritance was considered dominant. Genes were considered misexpressed when hybrids were significantly different from both parental populations. Patterns other than these were considered ambiguous.

### Allele Specific Expression and Mechanisms of Regulatory Divergence

We used the allelic depth (DP) field in the VCF file generated by *bcftools* to estimate the coverage of alternative alleles in each library. We restricted the allelic expression analysis to the sites identified previously as having fixed differences between subspecies. In order to examine patterns of allelic expression, we generated a data matrix containing counts for each site in each parental sample, and counts of each allele in hybrid samples. Thus the final data matrix contained twelve columns, one for each of the six parental samples, and two for each of the three hybrid samples (one for each allele). The site-level matrix of count data was normalized for read-depth in DE-Seq2, and differential expression tests were then used to identify sites showing significant regulatory divergence. For each site we conducted a test of differential allellic expression in the hybrids and for differential expression between the parental subspecies.

Evidence of biased allelic expression in F1 hybrids is a result of *cis* regulatory divergence (Wittkopp et al. 2004). *Trans* divergence is identified by comparing the ratio of expression in the parents and the ratio of expression of the alleles in hybrids (Wittkopp et al. 2004). To identify genes with significant *trans* regulatory divergence we constructed a linear model in DE-Seq2 with two main terms: “type”, which denotes whether reads were allelic counts in the parental species (note that each parental species only expresses on allele as these are fixed differences) or allelic counts in hybrids, which express both alleles. The second term describes the “condition”, whether counts are of allele A and B. The model also then included an interaction between condition and type,

~~~
design(transTest) <-formula(~ type + cond + type:cond)
resTransTest <-results(transTest, name="typeE.condB")
~~~

,where typeE specifies parental expression (as opposed to allelic) and condB, specifies allele B count. The “type” term in the model controls for differences in the counts between parental measurements and allelic measurements. This *results* function tests the null hypothesis that the ratio of allele A and B in the parental subspecies is equal to the allelic ratio of A to B in the hybrids. All tests were considered significant if the FDR-adjusted *p* value was less than 0.1.

Sites were categorized as *cis-only* if there was a significant expression difference between subspecies and there was allele-specific expression, but there was no evidence of *trans* divergence (Table 2). *Trans* only divergence was inferred if there was a difference between the subspecies, there was no allele specific expression in hybrids, but there was *trans* divergence. If there was was divergence in *cis* and *trans,* these could be further parsed into *cis* + *trans* and *cis* × *trans* based on whether parental divergence and allelic imbalance were in the same, or opposite direction, respectively. Compensatory evolution, a subcategory of *cis* × trans interactions, was inferred if there was no difference in expression between subspecies but there was evidence of divergence in both *cis* and *trans.* Sites that showed no parental, *cis* or *trans* divergence were considered conserved, and sites that did not fit any of these categories were considered ambiguous. Sites were functionally annotated using *SNPeff* (Cingolani et al. 2012), which uses the reference genome and annotation to determine where polymorphic sites are located relative to gene models.

## ACKNOWLEDGEMENTS

Thanks to Carol Goodwillie, Keith Keene and Polly Campbell for thesis committee service and for comments on earlier versions of the manuscript. Mike McCoy, Trisha Wittkopp, Joseph Coolon, Keith Adams, Katja Mack and Michael Love all provided helpful discussion of statistical analyses. Michael Love in particular provided the framework for testing for difference in allelic ratios (*trans* divergence) in DESeq2. Thanks to Katja Mack for providing access to her manuscript in review. This work was funded by East Carolina University. David Clayton kindly donated the Timor zebra finches used in this study and housing costs for all birds were supported by NIH NS045264 (Songbird Neurogenomics Initiative). Bin Luo performed the RNA extractions.

